# Basal amygdala inputs to the medial prefrontal cortex mediate fear memory strengthening

**DOI:** 10.1101/2022.01.20.477064

**Authors:** Einar Ö. Einarsson, Africa Flores, Daniel Jercog, Cyril Herry

## Abstract

Survival critically depends on the memorization of stimuli recurrently predicting pleasant or aversive experiences. Memory strengthening by additional learning is a critical process allowing the long-term stabilization of learned aversive experience. However, the underlying neuronal circuits and mechanisms are still largely unknown. Using a combination of single unit recordings and optogenetic manipulations, we observed that strengthened fear responses following successive fear learning episodes led to a remapping in the population of basal amygdala (BA) neurons encoding the conditioned stimulus. Moreover, strengthened fear responses were preferentially carried by BA neurons projecting to the medial prefrontal cortex (mPFC). Selective optogenetic inhibition of BA terminals in the mPFC during a second learning episode prevented fear memory strengthening. Together, our findings suggest that fear memory strengthening with additional training critically depends on the selective remapping of a subpopulation of BA neurons projecting to the mPFC.

## Introduction

When encountering stimulus that predicts danger, animals can form memories of these events through associative learning processes. Such associative fear learning serves a fundamental adaptive function in enabling organisms to modify their behavior to avoid danger in an ever-changing environment^1^. Experimentally, this has been studied by pairing an initially neutral stimulus (e.g. tone) with a mild aversive stimulus (e.g. foot-shock, the unconditional stimulus; US). Following learning, the tone becomes a conditioned stimulus (CS) that predicts the occurrence of the US, and can alone readily evoke conditioned fear responses such as an immobilization freezing response^2^. Following initial learning, associations can be modified through further experience, with repeated experience of the CS-US association leading to strengthening of conditioned responses, a phenomenon thought to depend on memory reconsolidation mechanisms^3^. In contrast, the repeated exposure to the CS alone leads to the weakening of the conditioned fear response through extinction learning^4,5^. While a number of studies have focused on the neuronal populations involved in weakening of the fear responses^6-8^, the neuronal populations involved in fear memory strengthening are virtually unexplored. A number of recent studies have suggested that learned fear is mediated by distributed neuronal networks of specialized neuronal subpopulations in multiple brain regions^8-10^. In particular, the basolateral amygdala, comprising the lateral amygdala, basal amygdala (BA), and basomedial subnuclei, is known to play a critical role in the acquisition and extinction of conditioned fear responses^11^. Notably, the amygdala contains functionally distinct classes of neurons that specifically respond to CS presentations during fear expression following conditioning, or extinction learning^7,12,13^. Indeed, longitudinal imaging studies in the basolateral amygdala have shown that conditioning induces such changes at individual and neuronal population levels, where CS representations become similar to US representations with training, whereas extinction induces novel representations dissimilar to those observed before or after the conditioning procedure^6,14^. However, to date, it remains largely unknown how fear memory strengthening following additional CS-US re-conditioning is encoded within dedicated circuits of the BA. In particular, it is still unexplored whether fear memory strengthening results in the recruitment of a new set of neurons or the selective refinement of cells taking part into an existing memory trace.

To address this question, we used a cued fear discrimination paradigm where mice were sequentially submitted to two conditioning sessions, each followed by a fear memory test session during which we monitored changes in neuronal activity within the BA using in vivo single-unit recordings during fear retrieval. This approach was combined with optogenetic and antidromic electrical identification of BA neurons projecting to the mPFC to precisely identify the neuronal circuits mediating fear memory strengthening. Finally, optogenetic manipulations were used to investigate the role of the BA-mPFC pathway in fear memory strengthening. Our results indicate that fear memory strengthening following a second conditioning session results in the remapping and decrease number of BA cells encoding the CS^+^ compared to that activated during the first memory retrieval session. Moreover, this subpopulation of BA neurons preferentially projects to the mPFC and the optogenetic inhibition of BA terminals in the mPFC prevented fear memory strengthening.

## Results

### Strengthened fear responding after re-conditioning

In order to address the question of how strengthened fear memory is encoded in the BA, we chronically implanted mice with recording electrodes in the BA, and trained them in a discriminative cued fear conditioning paradigm (**Figure 1A**). In this task, mice undergo mild fear conditioning session consisting of two paired presentations of a tone (CS^+^; 27 × 50 ms pips per presentation) and a foot shock (US) (Test 1). On the following day, mice are re-conditioned with the same two-pairing protocol (Test 2). As an internal control, a second stimulus (CS^-^; white noise) not paired with the US is also presented. This procedure induced strengthening on the group of mice as whole (**Figure 1B**). However, out of 21 implanted mice, 7 mice showed non-strengthened fear responses to the CS^+^ on Test 2 relative to Test 1, as defined by their discriminability index (d’) values of CS^+^ freezing during Test 1 and Test 2 being lower than 1 (**Figure 1C**). To compare possible differences in neural coding between fear strengthening and non-strengthening mice, we consider the same amount of mice showing the largest increase in strengthening as defined by the d’ values (**Figure 1C; Figure S1**). Accordingly, both strengthened and non-strengthened mice groups, froze more to the CS^+^ compared to the CS^-^ on both Test 1 and Test 2, demonstrating the specificity of the CS^+^ tone-fear association (**Figure 1D**). Importantly, it has been previously shown that a memory retrieval test session (CS+ presented without US) can lead to either stronger or weaker memory expression on a later retrieval session^15^. Thus, to test if increased freezing on Test 2 could be partially explained by a passive fear enhancement from Test 1, we omitted the re-conditioning session in a separate experiment (**Figure S1**). Our results indicate that the omission of the reconditioning session did not lead to fear memory strengthening on Test 2, thereby ruling out the possibility that our results could be simply explained by a passive fear enhancement from Test 1 retrieval session (**Figure S1**).

**Figure 1.**
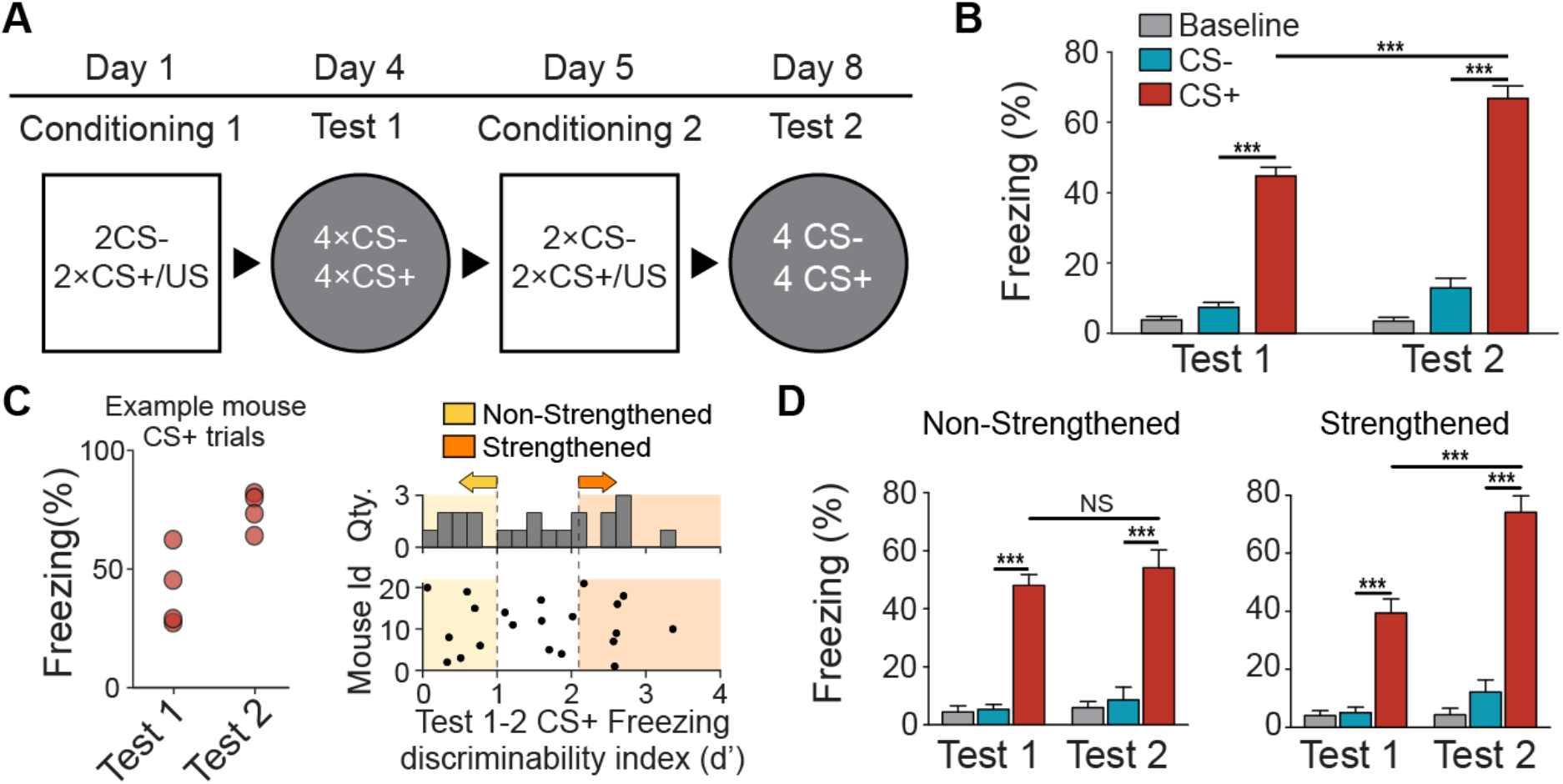
Fear strengthening protocol and behavioral performance. **A**. Experimental protocol for fear response strengthening, with two pairs of auditory fear conditioning sessions and test sessions. **B**. Behavioral results of fear conditioned mice (n = 21) showing increased freezing behavior to CS^+^ on Test 2 relative to Test 1 following a second conditioned session, while freezing during CS^-^ remain stable (two-way ANOVA: interaction, F_(2,126)_ = 13.29, p < 0.0001). **C**. Individual CS^+^ trial evoked freezing during Test 1 and Test 2 for an example mouse (right). Discriminability index (d’) for each mouse (left). Fear response was considered non-strengthened from Test 1 to Test 2 if d’ was lower than 1 (n = 7 mice). To compare with mice showing strong fear strengthening, the same amount of subjects (n = 7 mice) with the highest d’ values were considered as the strengthened group. **D**. Left, behavioral results of fear conditioned mice (n = 7) showing non-strengthened fear, while freezing during CS^-^ remain stable (two-way ANOVA: CS main effect F_(2, 24)_ = 177.9, p < 0.0001; CS^+^ freezing on Test 1 vs Test 2, NS p = 0.7026). Right, behavioral results of fear strengthened mice (n = 7), with freezing during baseline and CS^-^ remained stable (two-way ANOVA: interaction, F_(2,32)_ = 13.10, p < 0.0001).

### Encoding of fear strengthening in the BA

To identify activity changes related to fear memory strengthening, we performed single-unit recordings in the BA in freely behaving mice submitted to our behavioral paradigm. During retrieval Test 1 and Test 2 we recorded from 318 single units in the BA from the 14 mice (strengthened: 144 units, non-strengthened: 172 units) exposed to our behavioral paradigm (**Figure S2**). Out of the 70 / 90 units recorded in mice that showed strengthened / non-strengthened fear response on Test 2, 47 / 57 neurons were recorded on both Test 1 and Test 2 (**Figure 2A**; **Figure S2;** Methods).

**Figure 2.**
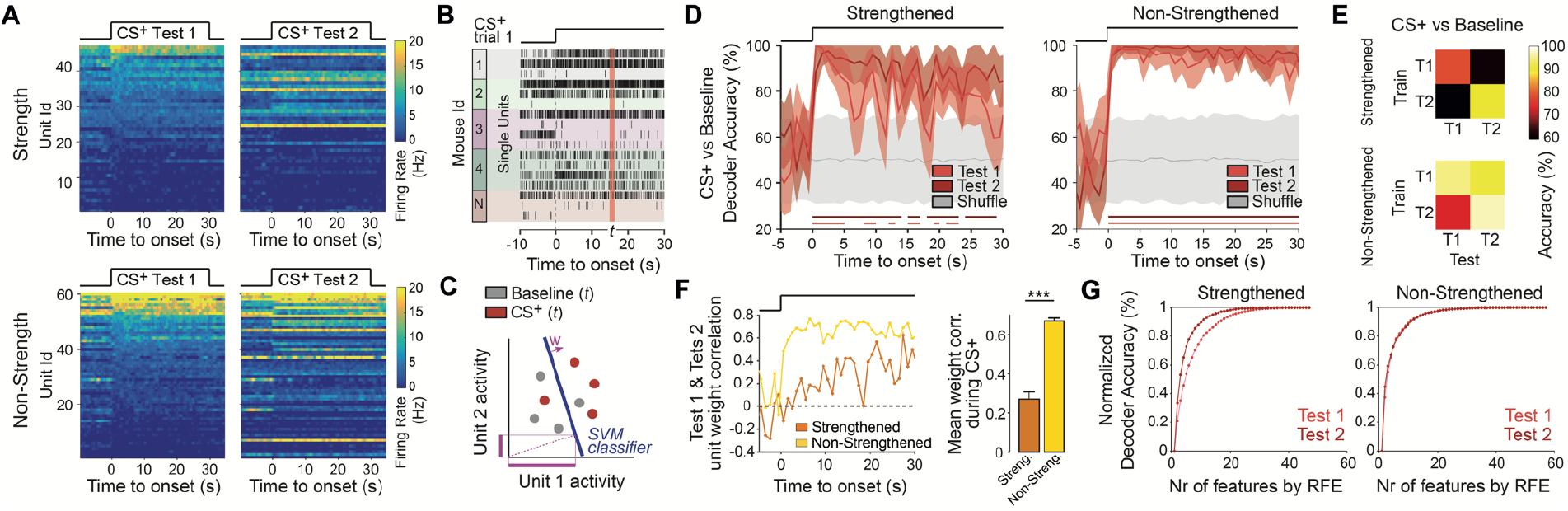
Fear strengthening leads to a reduced population in the BA encoding the conditioned stimulus. **A**. Firing rate heatmaps of BA single unit responses to CS^+^ during Test 1 and Test 2 from mice showing strengthened (top, n = 7 mice, 47 units) and non-strengthened (bottom, n = 7 mice, 57 units) fear responses on Test 2. Unit heatmaps ordered by average firing rate during first 10 s after CS onset during Test 1. **B**. Schematic of pseudo-population vector construction (*t*) on a single CS^+^ trial. Firing activity from different recorded cells from single animals in a randomly sampled trials are jointed across animals. **C**. Schematic of CS^+^ decoding analysis. For each time point (t, 1 s bins), independent support vector machine (SVM) linear classifiers were trained to discriminate CS^+^ from baseline activity. Individual unit contribution on the linear classifier accuracies is defined by the population decoder weights (w, averaged over cross-validations and repetitions). **D**. Mean decoding accuracy of CS^+^ from Baseline activity for Test 1 (light red) and Test 2 (dark red) and shuffled-labels. Thick lines indicate significant decoding accuracy timepoints (P < 0.05, Permutation test; 200 repetitions, 1000 shuffle; stratified leave-one out cross-validation). Error bars display mean +/-1 SD. **E**. Cross-time decoding accuracies from Test 1 and Test 2. Cross-time decoding performance drops to chance levels for the fear strengthened group but remains overall high for the fear non-strengthened group. **F**. Correlation of Test 1 and Test 2 individual-unit weight population decoder weight (averaged over cross-validations and repetitions) for fear strengthened and non-Strengthened mice (left). Average correlation values during CS^+^ presentations (right; paired T-test, p < 0.001). **G**. SVM recursive feature elimination on Test 1 and Test 2 for strengthened and non-strengthened groups. Bar plots indicate mean ± SEM. * *** p < 0.001.

To evaluate the impact of memory strengthening on CS representation in BA networks, we used a population decoding approach based on the recorded BA activity across mice^16^. This strategy is illustrated in **Figure 2B** in which the firing activity from the ensemble of recorded units from different mice at a given time point *t* (so-called pseudo-population vector) is compared on different conditions such as CS^+^ and CS^-^ trials versus baseline (**Figure 2C**; **Methods**). In the mice showing strengthened fear responses, CS^+^ from baseline decoders improved performance on Test 2 compared to Test 1, whereas decoding performance for the non-strengthened group remained similar (**Figure 2D**; **Figure S2**). Moreover, cross-time decoding on Test 1 and Test 2 days (i.e., training the decoders on one day and testing in the other) revealed that CS^+^ decoders performance for strengthened fear group dropped to chance levels whereas for the non-strengthened group performance remained above shuffle levels, suggesting stable CS^+^ representations from Test 1 to Test 2 for the non-strengthened fear group (**Figure 2E, Figure S2**). These data indicate that memory strengthening was associated with specific changes in population neuronal activity between Test 1 and Test 2. To further evaluate the nature of these population changes we analyzed the contribution of individual cells in the decoding performance observed in Test 1 and Test 2 by comparing their weight on the constructer decoders (**Figure 2C**). In agreement with the reduced stability of BA population representations across days for the fear strengthened mice (**Figure 2D**), we observed that individual decoding weights were less correlated from Test 1 to Test 2 for fear strengthened mice compared to the fear non-strengthened mice (**Figure 2F**). To address possible changes in the number of cells contributing to the decoders from Test 1 to Test 2, we performed a recursive feature elimination procedure by which the individual cells contributing less to the decoders are removed from the pooled cells in an iterative process^17^. Although the numbers of cells to reach maximal accuracy values for the non-strengthened fear mice did not differ from Test 1 to Test 2, more cells were necessary to reach maximal accuracy values on Test 1 compared to Test 2 for the strengthened group (**Figure 2G**). Together these data indicate that increased CS^+^ decoding performance observed in the fear strengthened mice was associated with both a remapping and a decrease in the populations of cells contributing to CS^+^ decoding from Test 1 to Test 2.

### Neurons encoding fear strengthening in the BA project to the mPFC

The BA is known to project to different regions of the mPFC to regulate fear expression and inhibition^8,9^. To examine if the BA-mPFC pathway is also involved in fear memory strengthening, we used antidromic activation of BA projections by using extracellular electrical stimulation in the mPFC following Test 2 in a cohort of mice submitted to our behavioral paradigm. (**Figure 3A**). Antidromic spikes were defined by high-fidelity, low-jitter and early latency response to stimulating pulses, in addition to entrainment to high-frequency stimulation (**Figure 3B**). Out of 48 BA units recorded on Test 2, 16 units were identified as projecting to the mPFC (33%). Notably, a higher fraction of units projecting to the mPFC were CS^+^ responsive (4 of 5 units) compared to those projecting elsewhere, suggesting that BA projecting to the mPFC were preferentially recruited for the encoding of CS^+^ during Test 2 (**Figure 3C**). Therefore, we next examined if the efferent pathway from the BA to the mPFC mediates fear strengthening during re-conditioning.

**Figure 3.**
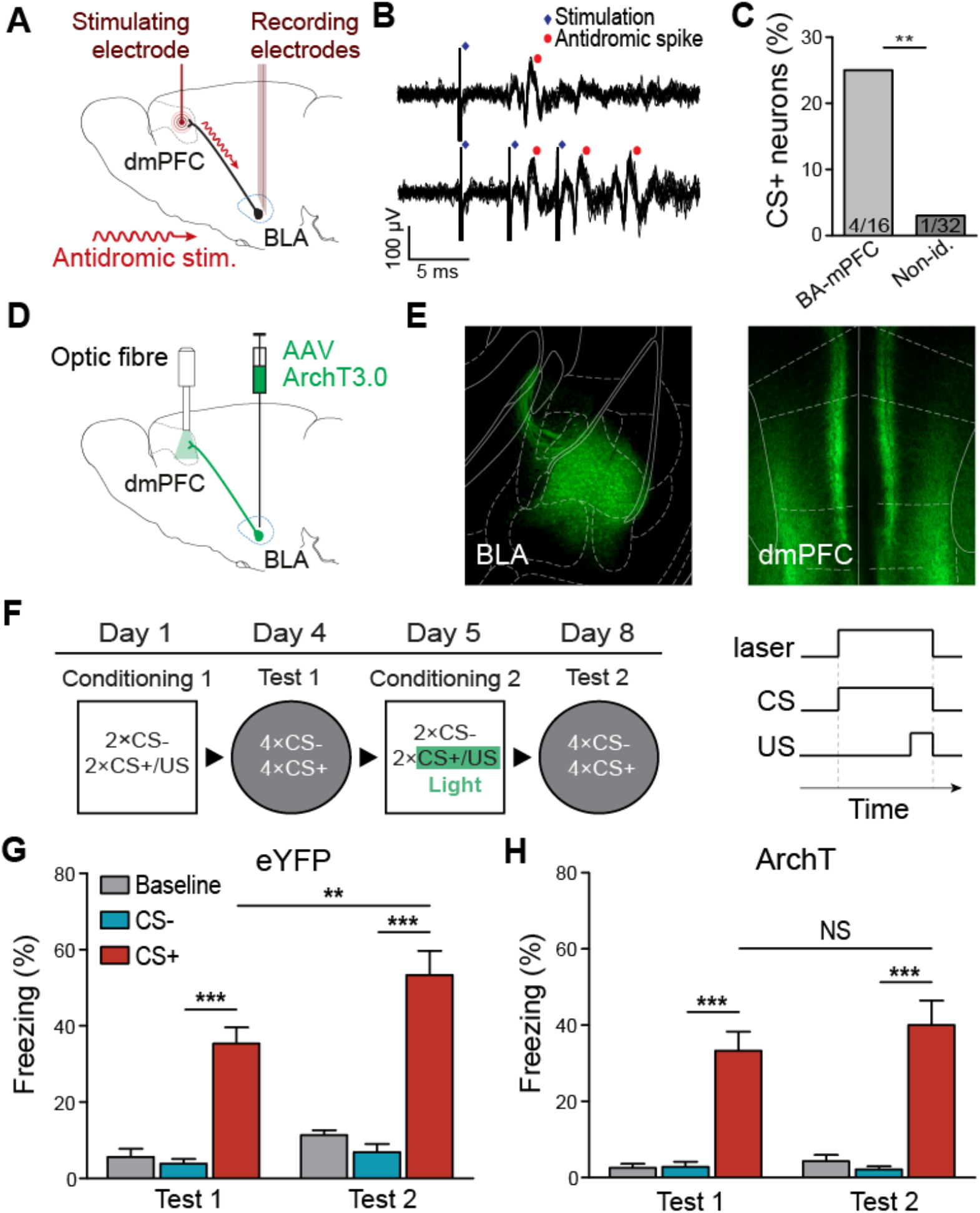
Inhibition of terminals of BA pyramidal neurons in the mPFC during CS^+^/US pairing impairs fear strengthening. **A**. Schema of experimental setup for detection of antidromic mPFC stimulation in the BA. **B**. Top and bottom, antidromic spikes recorded in the dmPFC following stimulation of the BA (10 superimposed traces) showing low-jitter, high-fidelity, and early-latency response of a BA neuron. **C**. Percentage of BA units in fear strengthened mice (n = 4 mice) with identified projections to the dmPFC or not, responding to CS^+^ on Test 2. BA neurons projecting to the mPFC were preferentially activated by CS^+^ (25%) relative to neurons not identified as projecting to the mPFC (3%) (exact binomial test, ** p < 0.01). **D**. Schematics of experimental design for optogenetic inhibition of ArchT infected BA neuronal projections in the mPFC. **E**. Neurons expressing CaMKII-eArchT3.0-eYFP in the mPFC (left) and BA (right). **F**. Experimental protocol for fear response strengthening, with two pairs of auditory fear conditioning sessions and test sessions, with light stimulation of the mPFC during the presentation of the CS^+^ and US in the second conditioning session. **G**. Mice infected with eYFP (n = 8) displayed increased freezing behavior in response to the CS^+^ than the CS^-^ on both Test 1 and Test 2, and increased CS^+^ freezing on Test 2 (two-way mixed design ANOVA: interaction, F_(1,21)_ = 8.03, p < 0.01). **H**. ArchT infected mice (n = 7) froze more to CS^+^ than CS^-^ on both Test 1 and Test 2, but did not show increased CS^+^ freezing on Test 2 (two-way ANOVA: CS main effect, F_(2,18)_ = 34.85, p < 0.0001). Bar plots indicate mean ± SEM. * p < 0.05; ** p < 0.01; *** p < 0.001.

We infected principal neurons in the BA with adeno-associated viruses (AAV) carrying the light-sensitive outward proton pump, archerodopsin-3 (ArchT) and enhanced yellow fluorescent protein (eYFP) (rAAV5-CamKIIa-eArchT3.0-eYFP), or eYFP only (rAAV5-CamKIIa-eYFP) as controls (**Figure 3D-E**). We then implanted optical fibers targeting the mPFC and photostimulated the infected BA-neuron terminals during the presentation of the CS^+^ and US-footshock during the second conditioning session (**Figure 3F**). We found that following re-conditioning, both ArchT and eYFP mice froze more to the CS^+^ than CS^-^ on both Test 1 and Test 2, but only eYFP mice showed increased CS^+^ freezing on Test 2 (**Figure 3G-H**). These data shows that the recruitment of BA-mPFC projecting neurons during the encoding of a second CS-US learning episode is necessary for fear memory strengthening.

## Discussion

In these experiments we used a combination of *in vivo* electrophysiology, electrical antidromic identification of BA projection neurons and optogenetic manipulation to characterize changes in the BA following fear memory strengthening, and identify the role of the BA-mPFC pathway in this phenomenon. We found that (i) strengthened behavioral fear responses following re-conditioning led to the remapping of BA neuronal population and the reduction of the number of cells recruited during memory strengthening, (ii) BA responding to the CS^+^ following fear memory strengthening preferentially project to the mPFC, and (iii) inhibiting BA terminals in the mPFC during re-conditioning prevented fear memory strengthening. Thus, our data identify for the first time a key role of the BLA-mPFC pathway in the regulation of cued fear memory strengthening.

These results indicate that cued fear memory strengthening, as revealed by a significant increase in freezing behavior from Test 1 to Test 2, is associated with a reduction of the number of BA cells encoding the CS^+^. This observation it at odds with the classical observation that changes in firing activity of amygdala neurons linearly predicts freezing responses. For instance, it has been shown that amygdala neurons gradually increase firing rate as a function of CS-US association but that extinction of the CS-evoked freezing response returns amygdala neurons activity to baseline^7,13^. In contrast, our results suggest that subtle changes in the temporal organization of the firing activity of BA neurons might be key mechanisms for fear memory strengthening, as documented for other neuronal structures and memory processes such as those taking place during sleep^18^. It is however not clear how these changes in BA CS^+^ response profile impact mPFC activity to drive fear responses. Based on the recent literature showing that (i) mPFC PV neurons precisely coordinate neuronal activity of mPFC PNs that preferentially project to the BLA to drive fear expression^19^ and (ii) optogenetic stimulation of BA terminals in the PL increases fear responding, whereas optogenetic inhibition reduces freezing^9^, it is likely that BA projections involved in memory strengthening impact on the very same circuit to enhance fear responding. Because BA inputs have been shown to preferentially contact dmPFC interneurons ^10,20,21^ it is tempting to speculate that memory strengthening on Test 2 could be due to a decrease drive of BA inputs onto dmPFC interneurons. Additional studies will be required in the future to disentangle these issues.

The reciprocally connected BA-mPFC pathway has been implicated in learning as well as expression of fear memory. Ex vivo recordings, show that tone-fear conditioning leads to strengthening of PL, but not IL, excitatory synapses in BA, by changes in postsynaptic increase in AMPA receptor function^22^. Moreover, synaptic depression of BA inputs to mPFC neurons impairs the recall of conditioned tone fear associations^23^. These studies suggest both the ascending and descending pathway between the BA and mPFC play a role in one trial tone-fear association conditioning. In line with these studies, we found that optogenetically inhibiting the BA inputs to the mPFC during re-conditioning blocked increased fear responding on a later test, i.e. inhibiting fear strengthening. This suggests that the BA-mPFC pathway plays a key role in additional fear learning following reconditioning, as that during one-trial fear conditioning learning. Finally, it remains to be determined how fear strengthening through re-conditioning leads to a reduction of the number of cells recruited and to the remapping of CS^+^ responsive neurons in the BA. Local inhibitory interneurons are known to provide strong regulation of PN output in the BLA^24^, where a disinhibitory microcircuit between PV and somatostatin-expressing interneurons has been shown to regulate BLA pyramidal neurons firing during the acquisition of associative tone fear memory^25^. Inhibitory feedback is thought to contribute to network oscillation and synchronization of BLA activity, and by improving spike-timing precision, could strengthen BLA signaling of the CS^+^ to the mPFC through refined synchronous firing activity of BA neuronal population following fear memory strengthening.

Interestingly, we observed that mice showing strengthened fear responses exhibited lower CS^+^ decoding accuracy compared to the mice that did not show strengthened fear responses on Test 1 (**Figure 2**). This observation sounds counterintuitive as the behavioral performance on Test1 was equivalent for both groups of mice. This could be due to the fact that behavioral performance on Test 1 does not merely reflect how the CS is encoded in BA networks. Alternatively, the lower CS^+^ decoding accuracy observed on Test 1 could be a neuronal signature of mice having the potential to display memory strengthening upon additional CS-US associations.

Imaging studies in the amygdala of pavlovian conditioned mice have shown that learning induce a similarity in CS representations to those representations elicited by the US, for both aversive^6^ and appetitive conditioning^14^. Accordingly, here we showed that strengthening of fear is associated with are remapping of CS^+^ representations. Moreover, we showed that the activation of mPFC projecting cells in the BA is required for such fear strengthening. Finally, further studies using longitudinal calcium imaging targeting BA neurons specifically projecting to the dmPFC will be required to evaluate the dynamic of neuronal changes associated with the strengthening of fear memories.

## Acknowledgements

We thank the members of the Herry laboratory for lively discussions and critical comments on the manuscript. We thank H. Wurtz for histological guidance, K. Deisseroth and E. Boyden for generously sharing materials, O. Yizhar for viral advices and S. Laumond and J. Tessaire and the technical staff of the housing and experimental facility of the Neurocenter Magendie. Microscopy was done in the Bordeaux Imaging Center of the CNRS-INSERM and Bordeaux University, member of France BioImaging. This work was supported by grants from the European Research Council (ERC) under the European Union’s Seventh Framework Program (FP7/2007-2013)/ERC grant agreement no. 281168, from the French National Research Agency (ANR-10-EQPX-08-OPTOPATH) and the Conseil Regional d’Aquitaine.

## Author Contributions

E.O.E., D.J., and A.F. performed behavioral experiments and analyzed the behavioral and electrophysiological data. C.H., D.J and E.O.E., designed the experiments and wrote the paper.

## Figures Legends

**Figure S1. Fear strengthening protocol behavioral performance**.

**A**. Individual mice performance on the strengthening protocol, showing freezing values for CS^-^ and CS^+^ during Test 1 and Test 2. Colored background indicates the mice is included on the fear non-strengthened and strengthened groups as define in **Figure 1C. B**. Freezing values during CS^+^ and CS^-^ presentations during habituation session for fear strengthened and non-strengthened groups showing no significant differences (two-way ANOVA: interaction, F_(1, 24)_ = 0.7981, p = 0.3805). **C**. Experimental protocol for testing fear response change without a second conditioning session. **D**. Behavioral result of fear conditioned mice (n = 7) showing decreased freezing behavior to CS^+^ on Test 2 relative to Test 1 (two-way ANOVA: interaction, F_(2,18)_ = 9.64, p < 0.01). Error bars indicate mean ± SEM. ** p < 0.01; *** p < 0.001.

**Figure S2. Single cell and population decoding results**.

**A**. Single cell p-values from t-test comparing average firing rate during 10 s before and after CS onset, for fear strengthened and non-strengthened mice. Pie charts showing the proportion of significant responsive cells defined by p < 0.05 (Strengthened: 144 units ; Non-strengthened : 172 units). **B**. Mean decoding accuracy of CS^-^ from Baseline activity for Test 1 (light gray) and Test 2 (dark gray) and shuffled-labels. No significant decoding accuracy timepoints observed (P > 0.05, Permutation test; 200 repetitions, 1000 shuffle; stratified leave-one out cross-validation). Error bars display mean +/-1 SD. **C**. Left column, mean decoding accuracy of CS+ from Baseline activity for decoders trained with data from Test 1 and tested with data from Test 2 for fear strengthened (top) and non-strengthened (bottom) groups. Right column, same but decoders trained with data from Test 2 and tested with data from Test 1.

**Figure S3. Recording and stimulation electrodes placement**.

**A**. Electrode placement for electrophysiological recordings in BA (n = 21 mice). **B**. Path of micro-stimulation electrodes for antidromic stimulation experiments (n = 4 mice, same color code as in panel A).

## Methods

### Experimental model and subject details

Male C57BL6/J mice (3 months old, Janvier) were individually housed for at least 7 days before all experiments, under a 12 h light–dark cycle, and provided with food and water ad libitum. All procedures were performed in accordance with standard ethical guidelines (European Communities Directive 86/60-EEC) and were approved by the committee on Animal Health and Care of Institut National de la Santé et de la Recherche Médicale and French Ministry of Agriculture and Forestry (agreement #A3312001).

### Behavior

Mice were trained using a discriminative cued fear conditioning paradigm during which one auditory conditioned stimulus (CS^+^) was paired with an aversive unconditioned stimulus (foot-shock) and another auditory stimulus (CS^-^) was presented alone. Behavioral testing was conducted in two distinctly different boxes (context A and B), where context A was used for conditioning, and context B for testing. Contexts A and B were cleaned with 70% ethanol and 1% acetic acid before and after each session, respectively. To score freezing behaviour, an automated infrared beam detection system located on the bottom of the experimental chambers was used (Imetronic). Animals were considered to be freezing if no movement was detected for 2 s. For all conditioning/test sessions, each CS presentation lasted 30 s and consisted of 27 50-ms pips (2 ms rise and fall) of a 7.5 kHz pure tone (CS^+^) or white-noise (CS^-^) at 80 dB (inter-trial intervals of 20 –180 s). Before conditioning (day 0), mice were habituate to the auditory stimuli in a session with 4 presentations of the CS^+^ and CS^-^. On day 1, mice underwent the discriminative cued fear conditioning in context A (Conditioning 1), where the CS^+^ was paired with the US (1-s foot-shock, 0.6 mA, 2x CS^+^/US pairings. The onset of the US coincided with the offset of the CS^+^. The CS^-^ was presented between each CS^+^/US pairing but was never paired with the US (inter-trial intervals of 20 –180 s). Three days later, the mice were tested for level of fear responding to the CS^+^ and CS^-^ in Context B (Test 1), where each CS was presented four times. The next day, the mice were re-conditioned in the exact same discriminative cued fear paradigm as before (Conditioning 2). Three days later, the mice were tested again for fear responding to the CS^+^ and CS^-^ in Context B as before (Test 2). In control experiment testing for passive fear enhancement after Test 1 (**Figure 1E-F**), the second conditioning session was omitted. Here, mice underwent Conditioning 1 (day 1) and Test 1 (day 4) as before, and were then tested again (Test 2) on day 8. In the optogenetic experiment (**Figure 4**), mice infected with either archaerhodopsin (ArchT) or yellow fluorescent protein (eYFP) in the BA underwent the same discriminative cued fear conditioning and re-conditioning as before, except that they received 31 s constant optical stimulation to the dmPFC during the presentation of CS^+^/US during the Conditioning 2 session. To evaluate the strengthening in fear responses between Test 1 and Test 2 For each mouse, freezing discrimination index d’ (**Figure 1C**) was defined as the absolute value of the sensitivity index, that is:

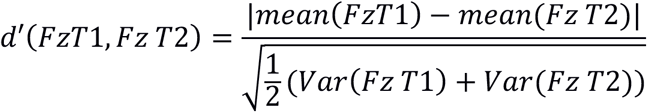

Where FzT1 and FzT2 are the freezing values for the 4 CS^+^ trials from Test 1 and Test 2, respectively.

### Surgery and recordings

Mice were anaesthetized with isoflurane (induction 3%, maintenance 1.75%) in O^2^ and body temperature was maintained at 37° C with a temperature controller system (FHC). Long-and short-lasting analgesic agents were injected during surgery (Metacam, Boehringer; Lurocaine, Vetoquinol). Mice were placed in a stereotaxic frame (Kopf Instruments) and 3 stainless steel screws were attached to the skull. Following craniotomy, mice were unilaterally implanted in the left BA with an electrode array at the following coordinates: 1.7 mm posterior to bregma; - 3.1 mm lateral to midline; and 4.3 mm ventral to dura. The electrode arrays consisted of 16 individually insulated nichrome wires (13 mm diameter, impedance 30–100 KU; Kanthal) fixed to a 26-gauge stainless-steel cannula guide. The electrode bundle was attached to an 18-pin connector (Omnetics), which was grounded via a silver wire (127 mm diameter, A-M Systems) soldered to a screw placed above the cerebellum. All implants were secured using Super-Bond cement (Sun Medical). After surgery mice were allowed to recover for at least 7 days. The electrode connector was connected to a headstage (Plexon) containing sixteen unity-gain operational amplifiers. Each headstage was connected to a 16-channel PBX preamplifier (gain 1000, Plexon) with bandpass filters at 300 Hz and 8 kHz. Spiking activity was digitized at 40 kHz and isolated by time-amplitude window discrimination and template matching using an Omniplex system (Plexon). At the conclusion of the experiment, electrolytic lesions were administered before transcardial perfusion to verify electrode tip location using standard histological techniques.

### Data analysis

Single-unit spike sorting was performed using Offline Sorter software (Plexon) and analyzed using Neuroexplorer (Nex Technologies) and MATLAB (MathWorks) for all behavioral sessions. Waveforms were manually defined while visualizing in a three-dimensional space using principal components, timing, and voltage features of the waveforms. A single unit was defined as a cluster of waveforms that formed a discrete, isolated, cluster in the feature space, and did not contain spikes with a refractory period less than 2 ms, based upon auto-correlation analyses. Additionally, multivariate ANOVA and J3 statistics were used to quantify separation of clusters in the principal component space. Cross-correlation analyses were performed to control that a single unit was not recorded on multiple channels. To assess unit stability between Test 1 and Test 2 recording session, waveforms recorded on each day were averaged and then correlated. Correlations with r values greater than 0.95 were considered stable units^7,26^.

### Single cell and population decoding analyses

For single cell analyses (**Figure S2**), individual units where considered CS responsive by comparing the average firing rate during the 10 s preceding and after CS onset showing a significant t-test with p < 0.05. To address how the firing rates of the BA cells ensembles were informative about the presence of a CS^+^, we used linear-kernel Support Vector Machine (SVM) classifiers^27^. Individual trial pseudo-population firing rate vectors were constructed for each 1 s time bin using units and trials from different animals, therefore removing decoding contributions due to noise correlations. For each time bin, an independent set of SVM classifiers were used to perform our decoding accuracies estimations and statistical tests. We used a “leave-one-out” stratified cross-validation (i.e., holding out 1 sample from each class) (**Figure 2**; **Figure S2**). The decoding accuracy of a classifier was defined as the proportion of correctly classified trials in the cross-validation procedure. To get an estimate of the accuracy variability, we performed a bootstrap 200 times on randomly selecting trials for pseudo-population trial construction. To evaluate the statistical significance of decoding accuracy, pseudo-population surrogate trial construction was obtained by randomly shuffling the label of the classes over 1000 bootstrap runs. The shuffled labels decoding was used to get a null distribution of the decoding accuracies that would occur by chance. For each time bin, we performed a Permutation test by computing P-values as the proportion of shuffle repetitions that exceed the real mean decoding accuracies^28^, and significant mean decoding accuracies were defined as those time bins where P < 0.05. Cross-time decoding was performed by training the SVM classifiers with data from one Test day and testing its performance with the data from the other Test day. Mean accuracy during CS^+^ values shown in **Figure 2E** are average across cross-validations and repetitions. Decoding weights are extracted after SVM model training and averaged across repetitions. Linear SVM recursive feature elimination was performed following^17^.

### Antidromic identification

Immediately following Test 2, a subset of mice (n = 4) were anesthetized with urethane (1.4 g kg^-1^) and secured in a stereotaxic frame. Concentric stimulating electrode (FHC) was lowered in the dmPFC at the following coordinates: 2 mm anterior to bregma; 0.3 mm lateral to midline; and 1.6 mm to 2 mm ventral to dura. During electric identification, the stimulation electrode was advanced in steps of 5 µm by a motorized micromanipulator (FHC) and evoked responses were recorded in the BA. Stimulation-induced and spontaneous spikes were recorded and sorted as described in Surgery and Recordings and Single-Unit Analyses. To ensure the same neurons were recorded during behavior and electric identification, waveforms averages from preceding behavioral session and the anesthesia session were tested for correlation as previously described. To be classified as antidromic, evoked-responses had to meet at least two out of three criteria^29^ : stable latency (< 0.3 ms jitter), collision with spontaneously occurring spikes, and follow high-frequency stimulation (250 Hz). At the end of the experiments, stimulating sites were marked with electrolytic lesions before perfusion, and electrode locations were verified as described in the Histological Analyses section.

### Virus injections and optogenetic manipulations

For pathway-specific optical inhibition of dmPFC terminals of CaMKIIa-expressing BA neurons, ArchT (rAAV5/CamKIIa-eArchT3.0-eYFP, UNC Vector Core Facility) was bilaterally injected into the BA of C57BL6/J mice from glass pipettes (tip diameter 10-20 mm) connected to a picospritzer (Parker Hannifin Corporation; 0.25 mL per hemisphere) at the following coordinates: 1.7 mm posterior to bregma, -3.1 mm lateral to midline, and 4.5 mm ventral to dura. At least 3 weeks after the injection mice were implanted bilaterally with custom-built optic fibers (diameter: 200 mm; numerical aperture: 0.39; Thorlabs) above the dmPFC at the following coordinates: 2.0 mm posterior to bregma, 0.6 mm lateral to midline, 1.2 mm ventral to dura, lowered at an angle of 10°. Control group was injected with eYFP (rAAV5/CamKII-eYFP, UNC Vector Core Facility). All implants were secured using 3 stainless steel screws and Super-Bond cement. After surgery mice were allowed to recover for at least 7 days. For optogenetic manipulation of ArchT-expressing CaMKIIa neurons, and matched eYFP controls, we delivered 31.5 s of continuous light, coinciding with the delivery of the CS^+^ and US. Green light (6 mW at fiber tip) was bilaterally conducted from a diode-pumping solid state laser (CNI Laser) to the mice via two fibre-optic patch cords (diameter 200 µ m, Doric Lenses), connected to a rotary joint (1 × 2 fibre-optic rotary joint, Doric Lenses) that allowed mice to freely move in the behavioral apparatus. The laser was controlled via TTL pulse generator (Master 9, AMPI), with each delivery ending with a 0.5 s ramping down period.

### Histological analyses

Mice were administered a lethal dose of isoflurane and underwent transcardial perfusions via the left ventricle with 4% w/v paraformaldehyde (PFA) in 0.1 M phosphate buffer solution (PB). Following dissection, brains were post-fixed for 24 hrs at 4°C in 4% PFA. Brain sections of 60 mm-thick were cut on a vibratome, and mounted on gelatin-coated microscope slides. To identify electrolytic lesions sections were stained with toluidine blue, dehydrated, mounted, and verified using conventional transmission light microscopy. Only electrodes terminating in the BA were included in our analyses. For verification of viral injections and optic fiber location in BA and dmPFC, respectively, mice were perfused as described above. Serial 60 mm-thick slices cut on a vibratome were treated with sodium borohydride (NaBH_4_ 1%, in 0.1 M PB, for 10 min) to reduce background auto-fluorescence (Clancy & Cauller, 1998) before being mounted in VectaShield (Vector Laboratories) and later imaged using an epifluorescence system (Leica DM 5000) fitted with a 10x dry objective. The location and the extent of the injections/infections were visually controlled. Only viral infections in the BA and optic fibers terminating in the dmPFC were considered for behavioral and electrophysiological analyses.

## QUANTIFICATION AND STATISTICAL ANALYSES

For all datasets, normality was tested using the Kolmogorov–Smirnov test (a < 0.05) and homogeneity of variance with Levene’s test (a < 0.05) to determine whether parametric or non-parametric analyses were required. Parametric analyses included t-test and two-way repeated-measures ANOVA followed by Bonferroni’s multiple comparison post hoc test if a significant main effect or interaction was observed. For categorical non-parametric comparisons, exact binomial test was used, with Fischer’s exact test of independence for 4×4 table comparison (**Figure 2C**). All tests were two-tailed and data are expressed as either mean ± s.e.m.; median, interquartile range, and extreme values; or mean ± 95% confidence interval. Sample sizes were determined based upon previous publications. Analyses were performed with MATLAB and Prism (GraphPad Software). Apart from t-tests, the asterisks in the figures represent the P-values of post hoc tests corresponding to the following values *p < 0.05; **p < 0.01; ***p < 0.001 based on mean ± s.e.m.

## DATA AND SOFTWARE AVAILABILITY

The data presented in this manuscript is available upon request to the corresponding author.

