## Supplementary figures and images for "Basal amygdala inputs to the medial prefrontal cortex mediate fear memory strengthening"

### Supplementary Figures 1-3

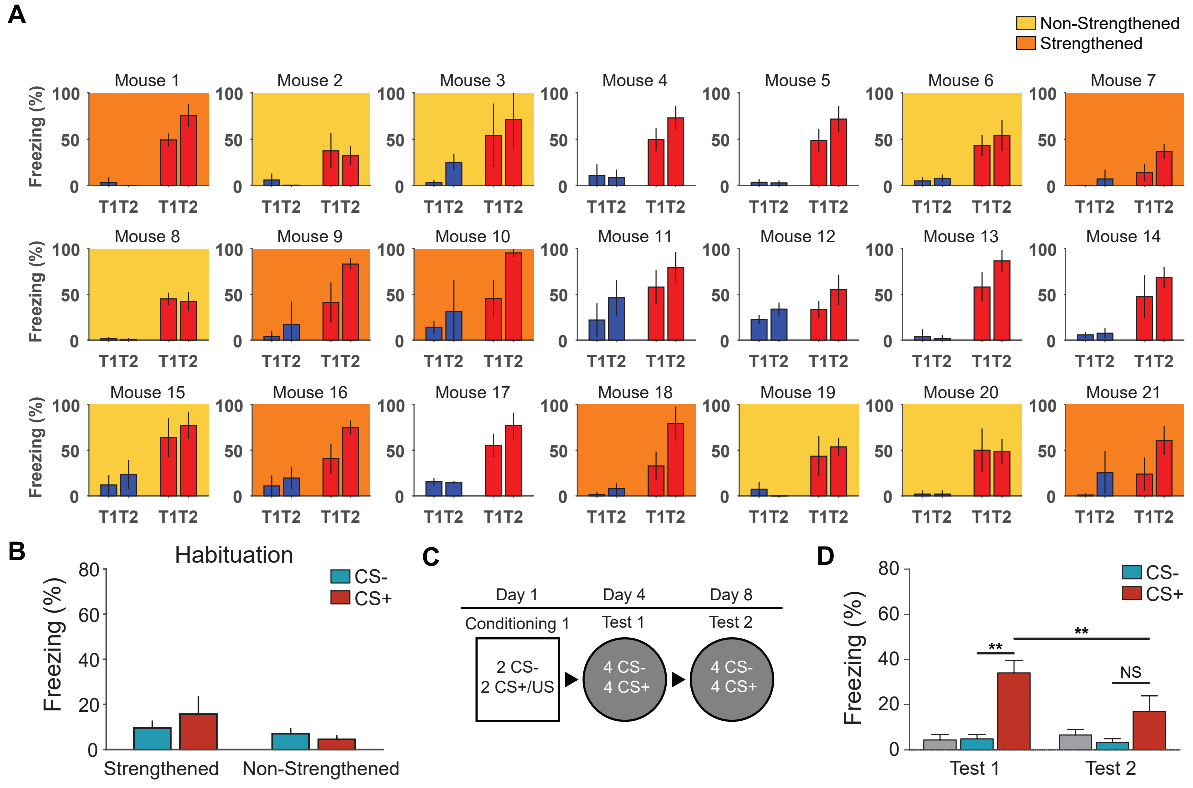

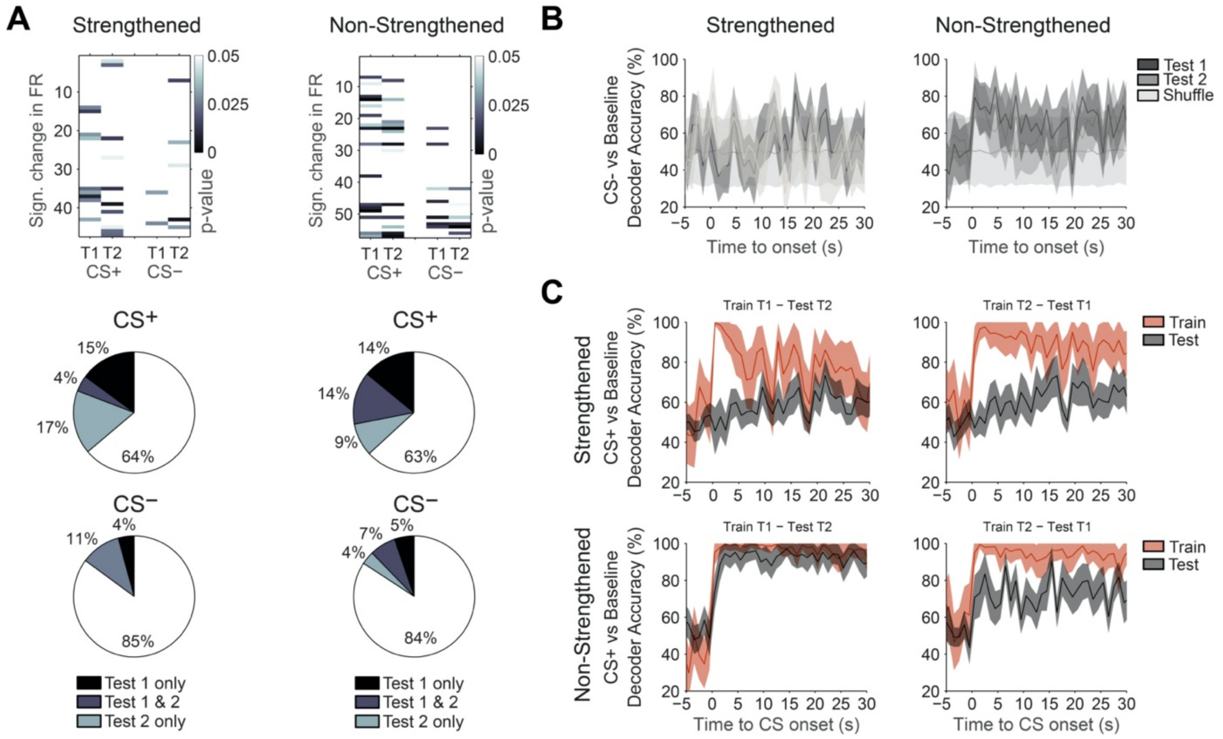

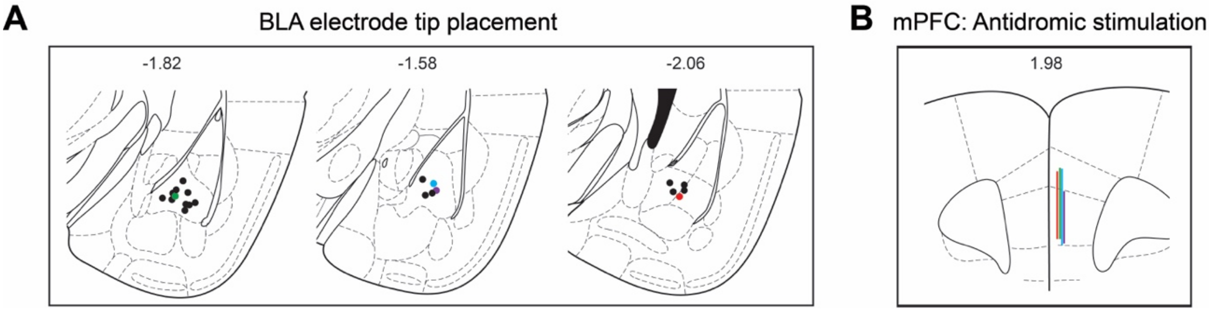
